# Multi-trait genome-wide analyses of the brain imaging phenotypes in UK Biobank

**DOI:** 10.1101/758326

**Authors:** Chong Wu

## Abstract

Many genetic variants identified in genome-wide association studies (GWAS) are associated with multiple, sometimes seemingly unrelated traits. This motivates multi-trait association analyses, which have successfully identified novel associated loci for many complex diseases. While appealing, most existing methods focus on analyzing a relatively small number of traits and may yield inflated Type I error rates when a large number of traits need to be analyzed jointly. As deep phenotyping data are becoming rapidly available, we develop a novel method, referred to as aMAT (adaptive multi-trait association test), for multi-trait analysis of any number of traits. We applied aMAT to GWAS summary statistics for a set of 58 volumetric imaging derived phenotypes from the UK Biobank. aMAT had a genomic inflation factor of 1.04, indicating the Type I error rates were well controlled. More important, aMAT identified 24 distinct risk loci, 13 of which were ignored by standard GWAS. In comparison, the competing methods either had a suspicious genomic inflation factor or identified much fewer risk loci. Finally, four additional sets of traits have been analyzed and provided similar conclusions.

## 1 Introduction

Genome-wide association studies (GWAS), which analyze a single trait each time, have identified thousands of genetic variants associated with an impressive number of complex traits and diseases [1]. While appealing, the identified genetic variants explain only a small proportion of the overall heritability, known as the ‘missing heritability’ problem [2]. In addition, many genetic variants are associated with multiple, sometimes seemingly unrelated traits [3]. This motivates multitrait association tests [4–9] that detect genetic variant associations with multiple traits by jointly analyzing these traits. Multi-trait association tests have successfully improved the statistical power and enhanced biological interpretations.

There is an exciting opportunity to develop novel and powerful multi-trait association tests that can analyze any number of traits jointly, as deep phenotyping data from epidemiological studies and electronic health records are becoming rapidly available [10, 11]. For example, GWAS of 3,144 brain image-derived phenotypes have been carried out to provide insights into the genetic architecture of the brain structure and function [11]. However, most existing multi-trait methods focus on a relatively small number (i.e., less than ten) of traits and it is unclear whether existing multi-trait association testing methods can handle any number of traits.

In this work, we develop a novel computationally efficient multi-trait association testing method, termed adaptive multi-trait association test (aMAT). Compared with many existing methods, aMAT has two compelling features that make it potentially useful in many settings. First, aMAT yields well-controlled Type I error rates when analyzing any number (e.g., hundreds) of traits. This is achieved by taking the potential singularity of the trait correlation matrix into account. In contrast, many competing methods yield incorrect Type I error rates. Second, aMAT maintains high statistical power (usually more powerful than competing methods) over a wide range of scenarios. Because the association pattern varies between SNPs, the most optimal test will also vary and is an unknown prior. To maintain high power, aMAT first constructs a class of test such that hopefully one of them may have good power for a given scenario and then combines the testing results data-adaptively. Through simulations, we demonstrate these two compelling features. Additionally, by analyzing several brain image-derived phenotypes GWAS summary results jointly, we demonstrate that our approach can reproducibly identify additional associated genetic variants that have been ignored by several existing methods. These newly identified genetic variants provide additional biological insights into brain structure.

## 2 Methods

### 2.1 Overview of aMAT

We build upon previous works [12–15] to develop a novel multi-trait association test for jointly testing the association between a single genetic variant and any number of (potentially hundreds) traits. We use one single nucleotide polymorphism (SNP) for illustration, and the same method can be applied across the whole genome. Suppose we have its *Z* scores across *p* traits of interest, ***Z*** = (*Z*_1_, *Z*_2_, *…, Z*_*p*_)^*′*^. Under the null hypothesis *H*_0_ that there is no association between the SNP and any traits of interest, ***Z*** follows a multivariate normal distribution [7–9] with mean zero and correlation matrix ***R*** = (*r*_*jk*_), i.e., ***Z*** ∼ *N* (0, ***R***).

aMAT involves three steps. First, we apply LD score regression (LDSC) [13, 14] to obtain an estimate of the correlation matrix, denoted by 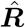 [9, 16]. Second, we construct a class of multi-trait association tests (MAT) such that hopefully each of them will be powerful under a certain scenario. To deal with potential (near) singularity problem of 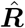, we calculate a modified pseudoinverse of 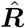, denoted by 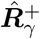. The test statistics of MAT is defined as follows:

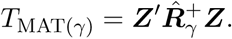

Third, because the uniformly most powerful test does not exist and the optimal value of *γ* is data-dependent and unknown, we propose adaptive MAT (aMAT) to combine the results from a class of MAT tests:

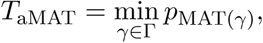

where *p*_MAT(*γ*)_ is the *p*-value of the MAT(*γ*) test and the default setting of Γ is Γ = {1, 10, 30, 50}, which performs well in both simulations and real applications. In the end, we apply a Gaussian copula approximation to calculate the *p*-value of aMAT efficiently.

### 2.2 Details of aMAT

Suppose we have *Z* score across *p* traits of interest for a SNP, ***Z*** = (*Z*_1_, *Z*_2_, *…, Z*_*j*_)^*′*^. Let ***β*** = (*β*_1_, *…, β*_*p*_)^*′*^ be the true marginal effect sizes for *p* traits. Note that *Z* score is either directly provided or can be calculated by 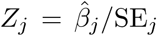, where 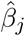 is the estimated effect size for trait *j* and SE_*j*_ is its corresponding standard deviation. We are interested in testing whether the SNP is associated with any trait of interest, i.e., *H*_0_: ***β*** = 0 vs *H*_1_: *β*_*j*_ ≠ 0 for at least one *j ∈* {1, 2, *…, p*}. Of note, testing multi-trait effect (*H*_0_: ***β*** = 0) is different from testing cross phenotype or pleiotropy effect [3], where the null hypothesis is at most one of the *β*_*j*_, *j* = 1, *…, p*, is nonzero.

Under the null hypothesis *H*_0_: ***β*** = 0, ***Z*** asymptotically follows a multivariate normal distribution with mean zero and correlation matrix ***R*** = (*r*_*jk*_), i.e. ***Z*** ∼ *N* (0, ***R***) [7–9]. Then we construct aMAT by the following three steps: 1) estimating trait correlation matrix ***R***; 2) constructing a class of multi-trait association tests (MATs); and 3) constructing adaptive MAT (aMAT) to maintain high power across a wide range of scenarios.

#### Estimating trait correlation matrix R

Following others [9, 16], aMAT applies LD Score Regression (LDSC) [13, 14] to obtain an estimate of ***R***, denoted by 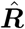. Specifically, we apply bivariate LDSC [14] to estimate the off-diagonal elements and univariate LDSC [13] to estimate the diagonal elements of ***R***. Because trait correlation matrix is the same for all the SNPs across the genome under the null, ***R*** only needs to be estimated once.

By applying LDSC, aMAT considers estimation error, including population stratification, cryptic relatedness, unknown sample overlap, and technical artifacts [16]. As demonstrated by others [9], LDSC provides a more accurate estimate of ***R*** than a commonly used approach that estimates ***R*** by sample correlation of genome-wide *Z* scores for each pair of traits [7, 8]. Thus, we choose LDSC to estimate the trait correlation matrix.

#### Constructing a class of MATs

Because the underlying association pattern is unknown, we construct a class of multi-trait association tests (MATs) that hopefully each of them would be powerful for a given scenario.

Many existing methods involve the inverse of estimated trait correlation matrix 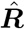. We observe that 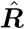 is often near singular when analyzing a large number of traits, leading to incorrect Type I error rates. To address this challenge, we propose a modified pseudoinverse approach. Specifically, we first apply the singular value decomposition (SVD) to 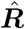 as 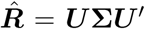, where ***U*** is a *p* × *p* orthogonal matrix and **Σ** is a *p* × *p* diagonal matrix that contains the descending ordered singular values *σ*_*i*_ of 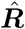 on its diagonal. We next calculate a modified pseudoinverse of 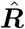 by

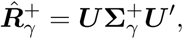

where 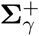 is formed from **Σ** by taking the reciprocal of the largest *k* singular values *σ*_1_, *…, σ*_*k*_, and setting all other elements to zero, where *k* is the largest integer that satisfies *σ*_1_*/σ*_*k*_ *< γ*. This is analogous to the principal components analysis, which restricts the analysis to top *k* axes of the largest variation. Next, we construct a class of multi-trait association tests (MATs) as:

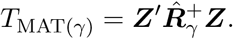

Of note, *T*_MAT(*γ*)_ follows a chi-squared distribution 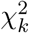 with *k* degrees of freedom under the null, and the *p*-value of MAT(*γ*) can be calculated analytically. MAT(*γ*) covers many existing tests as special cases. For example, MAT(1) equals to a principal component (PC)-based association test called ET [9]. MAT(50) equals to chi-squared test approximately when the estimated trait correlation matrix 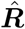 is not near singular.

#### Constructing aMAT

Because the association pattern varies between SNPs, the most optimal test will also vary and is an unknown prior. For example, MAT(1) achieves high power when the first principal component (PC) captures the majority association signals across the *p* traits. In contrast, when most PCs have weak signals, MAT(1) will lose power and MAT with larger *γ* will be more powerful. To maintain high power over a wide range of scenarios, we propose aMAT. Specifically, aMAT uses the smallest *p*-value from a class of MATs as test statistics:

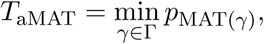

where *p*_MAT(*γ*)_ is the *p*-value of the MAT(*γ*) test, and a sensible default choice of Γ is Γ = {1, 10, 30, 50}. Similar methods have been widely used in gene-level association tests [12, 17], pathway-based analysis [18], and microbiome association analysis [19].

Next, to calculate the *p*-value of aMAT analytically, we apply a Gaussian copula approximation method, which has been successfully used in gene-level association tests [15]. Specifically, the Gaussian copula approximation-based method involves the following two steps.

- First, we estimate the correlation matrix **Ω** of the MAT(*γ*)s [*γ ∈* Γ] test statistics through parametric bootstrap under the null. Specifically, we first simulate a new *Z* score vector ***Z***^null^ under the null. In other words, we generate ***Z***^null^ by a multivariate normal distribution with mean zero and covariance matrix 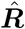. Next, we apply a class of MATs to the simulated *Z* score ***Z***^null^ and obtain their *p*-values. Under the null hypothesis, the *p*-values of MATs are uniformly distributed, and thus their inverse-normal transformed values (*q*_*γ*_ = Φ^−1^(1−*p*_MAT(*γ*)_)) follow a multivariate normal distribution with mean zero and covariance **Ω**. We repeat this procedure *B* (e.g., 10,000) times and then estimate **Ω** by the sample correlation matrix of *q*_*γ*_, denoted by 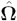. Because the null distribution for each SNP remains the same, this step only needs to be implemented once.
- Second, we apply a Gaussian copula approximation for the joint distribution of *q*_*γ*_ for *γ ∈* Γ. Specifically, the *p*-value of aMAT (*p*_aMAT_) can be calculated as

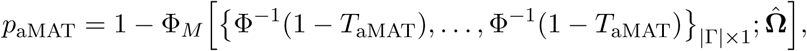

where Φ_*M*_ denotes joint distribution function of a multivariate normal distribution and |Γ| is the size of Γ.

### 2.3 Simulations

To speed up and simplify computations, we directly generated *Z* scores from its asymptotic normal distribution, *N* (Δ, ***R***). This simulation procedure has been widely used by others [9, 16]. The estimated trait correlation matrix of brain image-derived phenotypes (IDPs) [11] was used as the true trait correlation matrix ***R*** such that the simulation settings are similar to real applications. We mainly considered two scenarios: 1) the trait matrix for the set of 58 volumetric IDPs, denoted by Volume; and 2) the trait matrix for the set of 472 structural MRI related IDPs, denoted by Freesurfer (because those IDPs were derived by Freesurfer). To evaluate the impact of estimation error on ***R***, two simulation settings were considered. First, we conducted tests with the true trait correlation matrix ***R***. Second, we assumed ***R*** is unknown and conducted tests with the estimated trait correlation matrix 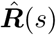. Specifically, similar to [16], independent normal noise with mean zero and variance *s* was added to each element of the matrix ***R*** to simulate the estimated trait correlation matrix, denoted by 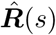. Based on our real data applications, the sampling variance was roughly 5 × 10^−5^. Thus we varied *s* from 10^−5^ to 10^−4^ to evaluate the impact of estimation error on ***R***.

We simulated 500 million or one billion *Z* score vectors under the null (Δ = 0) to evaluate Type I error rates with different significance level *α* and simulated 10, 000 *Z* score vectors under the alternative (Δ ≠ 0) to evaluate statistical power with the genome-wide significance level 5 × 10^−8^. Under the alternative, a wide range of scenarios were considered. For example, similar to [9], we generated 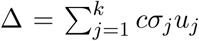, where *u*_*j*_ is the singular vector of the trait correlation matrix ***R***, *σ*_*j*_ is the *j*th singular value of ***R***, *c* is the effect size, and *k* is the largest integer that satisfies *σ*_1_*/σ*_*k*_ *< γ*. We varied *γ* and *c* to simulate different scenarios. We also considered other situations such as Δ = *cu*_*j*_ and Δ = *c*. To save space, the majority simulation results were relegated to the supplementary.

We compared aMAT with the following popular tests: 1) the sum of *Z* score vector (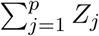 denoted as SUM) [6]; 2) the sum of squared *Z* score vector (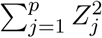; denoted as SSU) [5, 20]; 3) a standard chi-squared test; 4) a modified chi-squared test called Hom [8] that is powerful for the homogeneous effect situation; and 5) a PC-based association test [9], which equals to MAT(1). For chi-squared tests, we applied generalized inverse. When the trait correlation matrix is near singular (our focus here), two popular tests Het [8] and AT [9] had some numerical issues, failing to provide *p*-values and thus having been ignored in our comparisons. The minP test [4] that takes the minimum marginal *p*-values across *p* traits as the test statistics is highly conservative and time-consuming when *p* is large and thus has only been briefly discussed.

### 2.4 Analysis of UK Biobank brain imaging GWAS summary data

We reanalyzed GWAS summary results of image-derived phenotypes (IDPs) with up to 8,428 individuals [11] in UK Biobank to gain additional insights into the genetic architecture of brain imaging. Specifically, we conducted multi-trait association tests to detect the genetic association between an SNP and five sets of highly related IDPs with up to 472 IDPs per set. These five sets were determined by [11], covering a wide range of the IDP classes with significant trait correlations in each grouping. To be concrete, we focused on the set of 58 brain volumetric measures [11] and briefly discussed the additional four IDP sets.

#### Data pre-process and multi-trait association tests

We removed variants that were either non-biallelic or strand ambiguous (SNPs with A/T or C/G alleles) and analyzed the remaining 9,971,805 SNPs. LDSC was applied to estimate the trait correlation matrix. For each set of IDPs, we applied aMAT with default setting (Γ = {1, 10, 30, 50}) and many competing methods, including SUM, SSU, MAT(1) (equals to ET), Chi-squared, and Hom tests to test the overall association between a SNP and a pre-defined IDP set.

#### Result analyses

We used the genomic inflation factor to evaluate the empirical Type I error rates. The genomic inflation factor is defined as the ratio of the median of the observed test statistic to the expected median, which quantifies the extent of inflation and false positive rate. An ideal genomic inflation factor is one. However, because of the polygenic nature of GWAS signals, a genomic inflation factor is usually slightly larger than one. However, a genomic inflation factor greater than 1.2 or even 1.5 shows an indication of inflated Type I error rates, and a genomic inflation factor less than 0.9 shows evidence for conservative Type I error rates.

In addition, we used Functional Mapping and Annotation (FUMA) [21] v1.3.4c to identify independent genomic risk loci and lead SNPs in these loci. Via FUMA, We further obtained the functional consequences for these SNPs by matching SNPs to many functional annotation databases, including CADD scores [22], REgulomeDB scores [23], chromatin states [24, 25], and ANNOVAR categories [26]. For lead SNPs identified by aMAT, replication was tested by an independent dataset obtained from ENIGMA consortium [27]. Specifically, we applied aMAT to the GWAS summary statistics of seven brain subcortical volumetric IDPs, including the volume of mean putamen, the volume of mean thalamus, the volume of mean pallidum, the volume of mean caudate, Intracranial Volume, the volume of mean amygdala, and the volume of mean accumbens. Then the replication rate was evaluated by the two-tailed binomial test. In the end, we filtered genome-wide significant SNPs identified by aMAT based on their functional annotations and then mapped them to genes by the following three strategies (via FUMA): positional mapping, eQTL mapping, and chromatin interaction mapping.

### 2.5 Availability of data and materials

The UK Biobank summary data is available in the http://big.stats.ox.ac.uk. The LDSC software and its required LD scores are available in the https://github.com/bulik/ldsc. The validation data from NIGMA Consortium can be obtained from http://enigma.ini.usc.edu/research/download-enigma-gwas-results/. The software for our proposed method aMAT is available at https://github.com/ChongWuLab/aMAT. Supplementary files available at https://figshare.com/articles/aMAT_supplementary/9764519.

## 3 Results

### 3.1 aMAT yields well-controlled Type I error rates

Table 1 shows the Type I error rates of different multi-trait methods with the Freesurfer trait correlation matrix. Our proposed methods MAT and aMAT yielded well controlled Type I error rates at different significance levels *α*. Of note, the Freesurfer set contained 472 IDPs, and 27 eigenvalue values of the trait correlation matrix were smaller than 0.01 (Supplementary Figure S1). As expected, chi-squared and Hom tests yielded inflated Type I error rates because corresponding test statistics involve the inverse of a near singular correlation matrix. Also, SSU yielded inflated Type I error rates. All methods except SSU yielded well-controlled Type I error rates under simulations with Volume trait correlation matrix and thus were relegated to Supplementary (Supplementary Table S1).

**Table 1:**
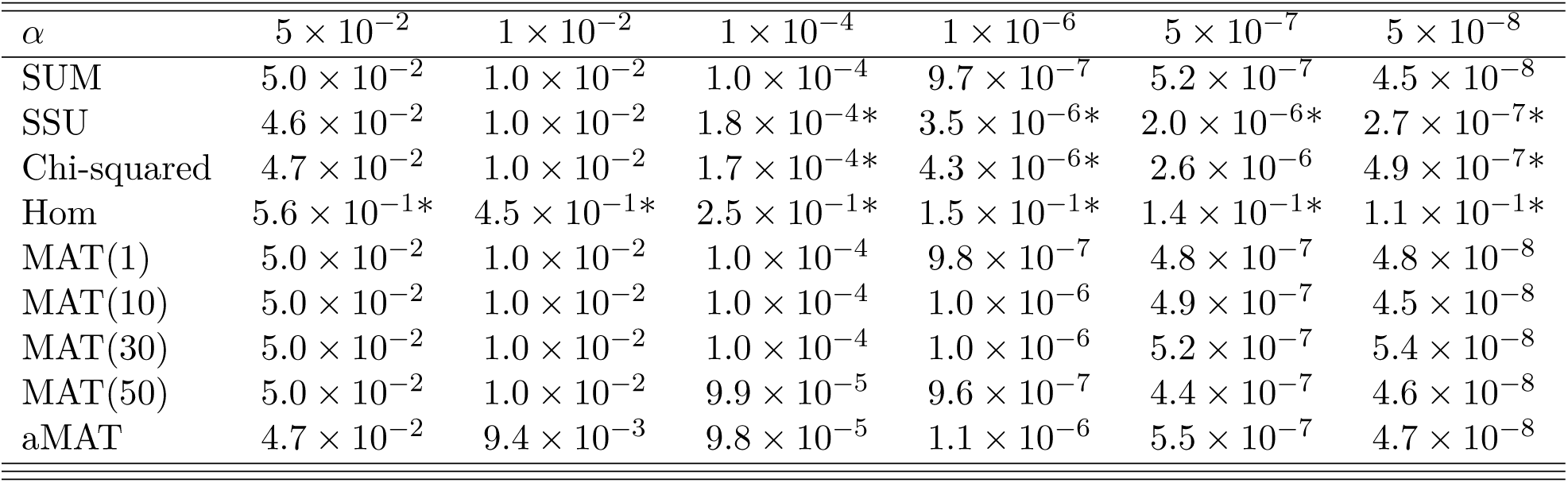
Type I error rates of different methods with the Freesurfer trait correlation matrix. We simulated one billion (1 ×10^9^) replications under the null and estimated Type I error rates as the proportions of *p*-values less than significance level *α*. * inflated Type I error rates.

### 3.2 aMAT offers robust statistical power

Figure 1 shows the empirical power of different methods with true Volume trait correlation matrix ***R***. When the first principal component (PC) was informative, as expected, the MAT(1) performed best. However, MAT(1) was sensitive to the signal distribution and yielded much lower power when the top PC has weak or no signal (Supplementary Figure S2). The minP test was very conservative, and the power was almost zero for the situations considered here. Hom test [8] was known to be powerful for the homogeneous effect situation, but the power was close to zero when many principal components were informative (Figures 1d, e, and f). SUM and SSU tests ignored the relationships among traits and thus were less powerful than MAT tests. Because there is no uniformly optimal test for a composite alternative *H*_1_: ***β*** ≠0, different MAT tests achieved high power under different scenarios. However, aMAT maintained high power across a wide range of scenarios by combining the results from a class of MAT tests.

**Figure 1:**
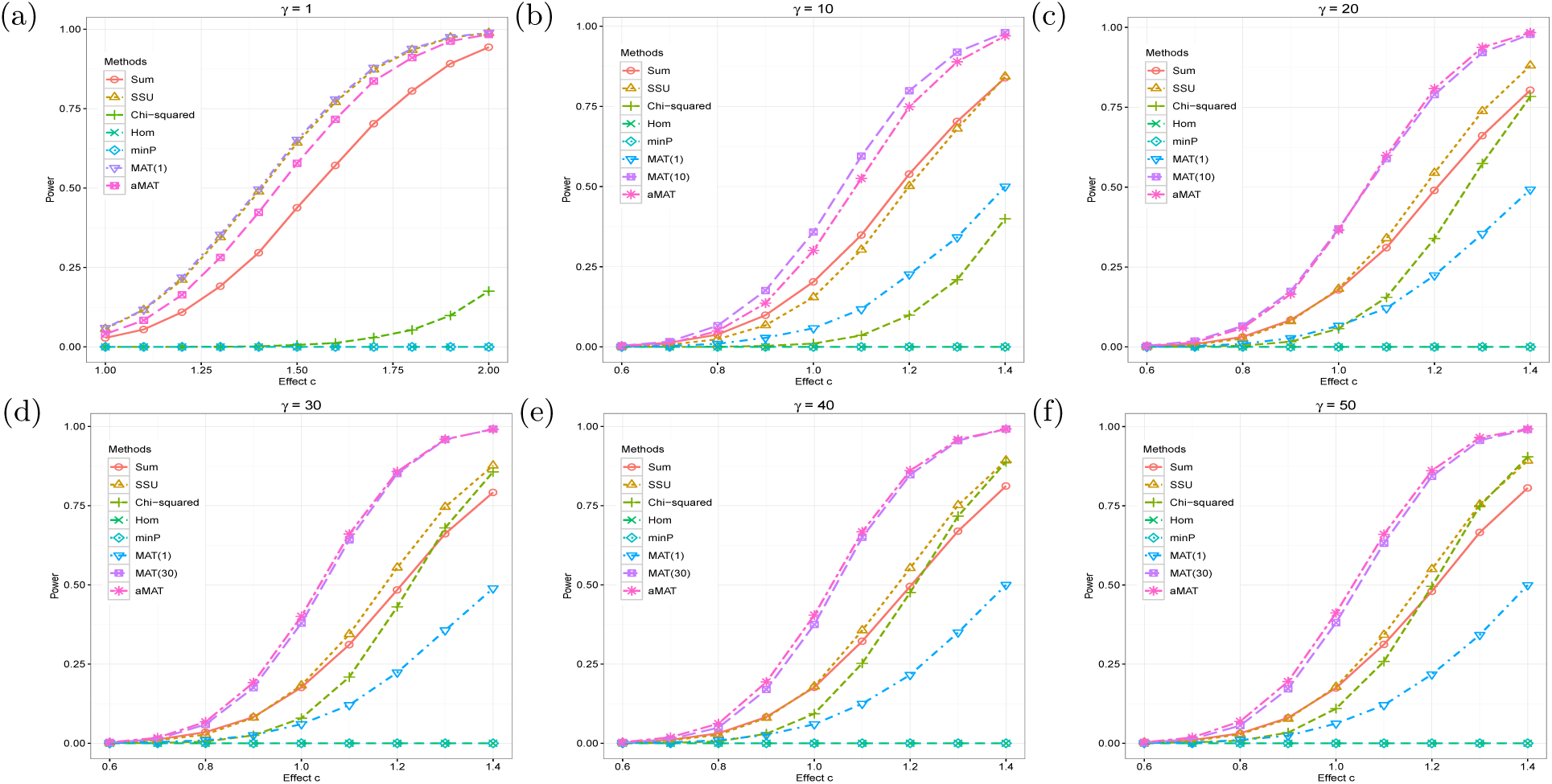
Empirical power comparison with true Volume trait correlation matrix. Under the alternative, we generated 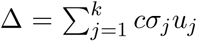, where *u*_*j*_ is the *j*th singular vector of the Volume trait correlation matrix ***R***, *σ*_*j*_ is the *j*th singular value, *c* is the effect size, and *k* is the largest integer that satisfies *σ*_1_*/σ*_*k*_ *< γ*. Empirical power was estimated as the proportions of *p*-values less than significance level 5 × 10^−8^.

We considered several additional settings, including varying the way of selecting informative singular vectors and generating Δ, the trait correlation matrix (including five traits, 25 traits, Volume (58 traits), Area (206 traits), Thickness (208 traits), and Freesurf (472 traits)), and the generating distribution for Δ (Supplementary Figures S3–S13). We show that aMAT achieved a robust statistical power under all simulations considered. In contrast, competing methods that achieved high power under one specific situation could lose power substantially under several other situations.

### 3.3 aMAT is robust to the estimation error of trait correlation matrix

In the previous two subsections, we assumed ***R*** was known. However, the trait correlation matrix ***R*** is unknown and has to be estimated. To evaluate the impact of estimation error on ***R***, we simulated *Z* score vectors from its asymptotic normal distribution *N* (Δ, ***R***) and compared different testing methods with the estimated trait correlation matrix 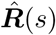, which was constructed by adding independent normal noise with mean zero and variance *s* to each element of ***R***.

Table 2 shows the Type I error rates of different methods with the estimated Volume trait correlation matrix 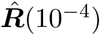. Both MAT and aMAT yielded well-controlled Type I error rates. In comparison, both chi-squared and Hom tests yielded inflated Type I error rates even though they had controlled (slightly conservative) Type I error rates with the true Volume trait correlation matrix ***R*** (Supplementary Table S1). This is because both chi-squared and Hom test statistics involve the inverse of the estimated trait correlation matrix 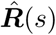 and small estimation error could lead to large deviation for small eigenvalues.

**Table 2:**
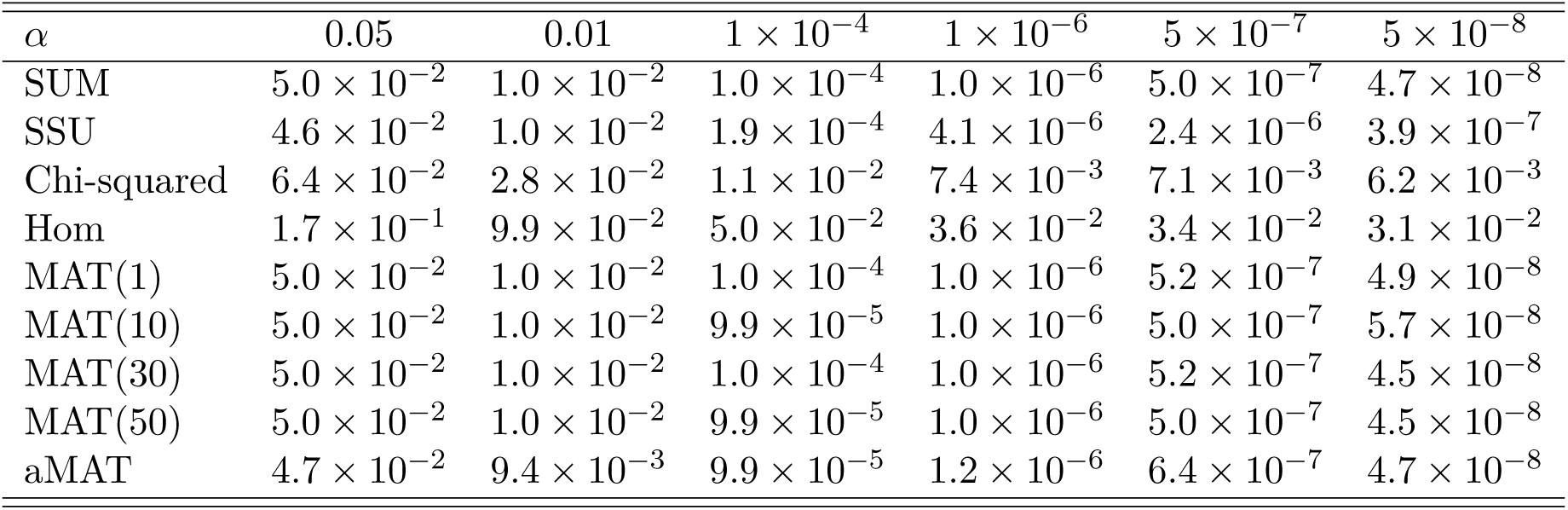
Type I error rates for different methods with the estimated Volume trait correlation matrix 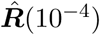. We simulated 500 million (5 ×10^8^) replications with true Volume trait correlation matrix ***R*** under the null and constructed test statistics with 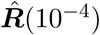. Type I error rates were estimated as the proportions of *p*-values less than significance level *α*.

We next evaluated the impact of estimation errors on the statistical power. Figure 2 shows that the empirical power of MATs and aMAT with either true (***R***) or estimated 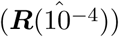 trait correlation matrix were almost the same, indicating estimation errors had little impact on the power of MATs and aMAT.

**Figure 2:**
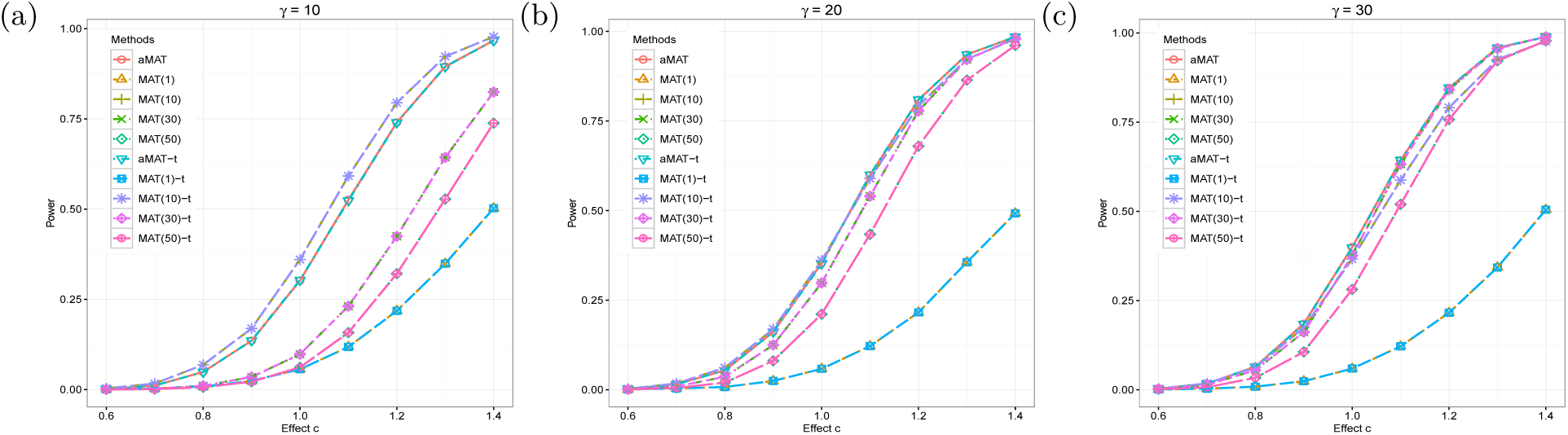
Empirical power comparison between true and estimated Volume trait correlation matrix. Under the alternative, we generated 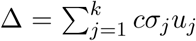, where *u*_*j*_ is the singular vector of the true Volume correlation matrix ***R***, *c* is the effect size and *k* is the largest integer that satisfies *σ*_1_*/σ*_*k*_ *< γ*. We simulated 10,000 replications with ***R*** and constructed test statistics with 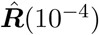. We further estimated empirical power as the proportions of *p*-values less than significance level 5× 10^−8^. MAT(1), MAT(10), MAT(30), MAT(50), aMAT represent the results with 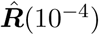, while MAT(1)-t, MAT(10)-t, MAT(30)-t, MAT(50)-t, aMAT-t represent the results with ***R***.

In the end, we considered several additional settings with different *s* (*s* = 10^−5^ or *s* = 5 × 10^−5^) and/or with the estimated Freesurf trait correlation matrix. The results were similar to those in Table 2 and Figure 2, thus were relegated to the Supplementary (Supplementary Tables S2–S6 and Supplementary Figures S14–S18).

### 3.4 aMAT identifies novel risk loci for brain volumetric measures

To demonstrate aMAT’s effectiveness in real multi-trait association studies, we performed a multistage study for a set of 58 volumetric image-derived phenotypes (IDPs), which was defined by [11]. In the discovery stage, we applied aMAT to the GWAS summary statistics from UK Biobank [11]. aMAT had a genomic inflation factor of 1.04 (Supplementary Figure S19), indicating the type I error rates of aMAT were well controlled. aMAT identified 801 significant SNPs (with *P <* 5 × 10^−8^) across the whole genome, 453 of which were ignored by any individual IDP tests at the 5 × 10^−8^ genome-wide significance level. These 801 GWAS variants were represented by 28 lead SNPs, located in 24 distinct risk loci (Table 3 and Figure 3). Among these 28 lead SNPs, 13 SNPs (46.4%) were missed by any individual IDP tests at the 5 × 10^−8^ genome-wide significance level.

**Table 3:** Summary statistics of significantly associated regions identified by aMAT in the multi-trait analysis of a set of volume related IDPs. Independent lead SNPs are defined by *r*^2^ *<* 0.1 and distinct loci are *>* 250kb apart. SNP, CHR, BP, A1, A2 are the lead SNP, chromosome, position, effect allele, and non-effect allele, respectively. The MAF is the minor allele frequency and obtained from the 1000 genomes reference panel (Phase 3). aMAT P is the *p*-value for the aMAT test. The bolded SNPs correspond to the novel SNPs that were ignored by any individual IDP tests at the genome-wide significance level 5 × 10^−8^.

| Locus | SNP | CHR | BP | A1 | A2 | MAF | aMAT P | Nearest Gene |
| --- | --- | --- | --- | --- | --- | --- | --- | --- |
| 1 | **rs72688340** | 1 | 47983839 | G | A | 0.171 | $2.5 \times 10^{-9}$ | *AL356458.1* |
| 2 | **rs6665134** | 1 | 180949628 | C | A | 0.401 | $1.7 \times 10^{-10}$ | *STX6* |
| 3 | rs476557 | 1 | 215140251 | A | C | 0.475 | $4.3 \times 10^{-8}$ | *RP11-323K10.1* |
| 4 | **rs1504** | 2 | 37066018 | T | G | 0.447 | $1.4 \times 10^{-8}$ | *AC007382.1* |
| 5 | rs13070564 | 3 | 190629975 | G | T | 0.383 | $8.8 \times 10^{-13}$ | *GMNC* |
| 6 | **rs10031823** | 4 | 103125031 | T | C | 0.395 | $1.3 \times 10^{-8}$ | *SLC39A8* |
| 6 | rs34333163 | 4 | 103283117 | A | G | 0.075 | $4.4 \times 10^{-16}$ | *SLC39A8* |
| 7 | **rs1922930** | 6 | 1364691 | A | C | 0.118 | $3.8 \times 10^{-8}$ | *RP4-668J24.2* |
| 8 | **rs77126132** | 7 | 54966738 | G | A | 0.089 | $1.8 \times 10^{-8}$ | *SNORA73* |
| 9 | rs2707521 | 7 | 120940436 | C | T | 0.377 | $2.2 \times 10^{-16}$ | *CPED1* |
| 10 | **rs2974298** | 8 | 42376477 | T | C | 0.493 | $2.5 \times 10^{-9}$ | *SLC20A2* |
| 11 | rs10217651 | 9 | 118923652 | A | G | 0.386 | $1.7 \times 10^{-14}$ | *PAPPA* |
| 11 | rs35565319 | 9 | 119065043 | C | T | 0.072 | $6.7 \times 10^{-16}$ | *PAPPA* |
| 11 | **rs7030607** | 9 | 119245183 | G | A | 0.36 | $6.7 \times 10^{-9}$ | *ASTN2* |
| 12 | **rs12783517** | 10 | 21878407 | C | T | 0.299 | $1.3 \times 10^{-8}$ | *MLLT10* |
| 13 | rs4962692 | 10 | 126424823 | G | A | 0.414 | $2.7 \times 10^{-10}$ | *FAM53B* |
| 14 | **rs1187159** | 11 | 92009792 | T | C | 0.413 | $5.1 \times 10^{-11}$ | *NDUFB11P1* |
| 15 | rs73123652 | 12 | 65874956 | T | C | 0.106 | $1.1 \times 10^{-12}$ | *MSRB3* |
| 16 | rs4301837 | 12 | 102336310 | T | C | 0.496 | $6.9 \times 10^{-9}$ | *DRAM1* |
| 16 | rs17797222 | 12 | 102913946 | A | G | 0.224 | $3.6 \times 10^{-13}$ | *RP11-210L7.1* |
| 17 | rs12146713 | 12 | 106476805 | T | C | 0.093 | $3.2 \times 10^{-9}$ | *NUAK1* |
| 18 | rs77956314 | 12 | 117323367 | T | C | 0.083 | $1.2 \times 10^{-9}$ | *HRK* |
| 19 | rs10129414 | 14 | 56193272 | G | A | 0.438 | $2.0 \times 10^{-9}$ | *RP11-813I20.2* |
| 20 | rs2464469 | 15 | 58362025 | G | A | 0.412 | $2.1 \times 10^{-13}$ | *ALDH1A2* |
| 21 | **rs13330163** | 16 | 70660243 | A | G | 0.453 | $3.0 \times 10^{-12}$ | *IL34* |
| 22 | rs12920553 | 16 | 87227046 | G | T | 0.418 | $3.5 \times 10^{-10}$ | *C16orf95* |
| 23 | **rs6121038** | 20 | 30254773 | T | G | 0.291 | $5.2 \times 10^{-10}$ | *BCL2L1* |
| 24 | **rs1004764** | 22 | 38474852 | G | A | 0.377 | $4.2 \times 10^{-9}$ | *SLC16A8* |

**Figure 3:**
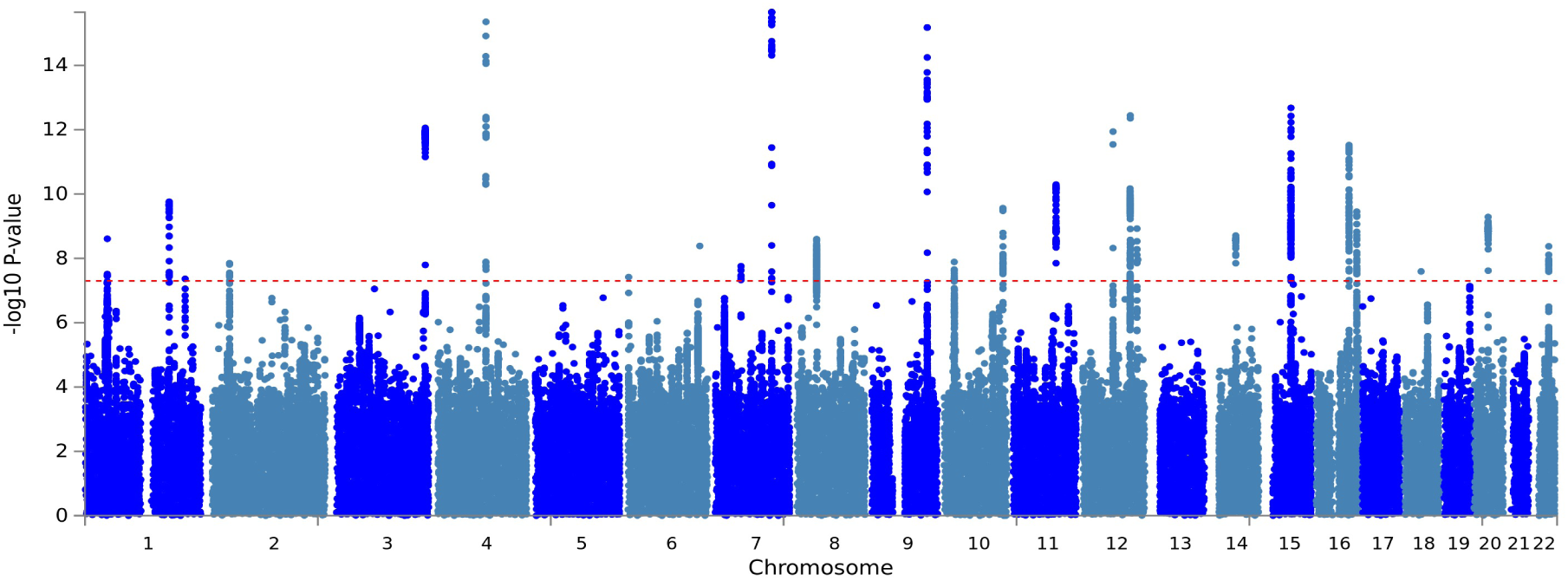
Multi-trait analysis for the Volume set. Manhattan plot displays the association results of aMAT per variant ordered by their genomic position on the *x* axis and showing the strength with the − log_10_(*p*) on the *y* axis.

Next, we replicated our findings in an independent dataset obtained from the ENIGMA consortium [27], which contains GWAS summary statistics of seven subcortical volumes in up to 13,171 subjects. Despite seven brain subcortical volumetric IDPs reported in ENIGMA consortium only represent a subset of the volumetric IDPs in UK Biobank, the replication rate was high. Out of the 24 distinct risk loci, four loci (rs73123652, rs77956314, rs10129414, and rs1187159) were successfully replicated (two-tailed binomial test *P* = 6.25 × 10^−30^) under the genome-wide significance threshold (*P <* 5 × 10^−8^)). The numbers of replicated SNPs rose to 13 under a relaxed cutoff of 0.05 (two-tailed binomial test *P* = 2.24 × 10^−10^).

To link the identified variants to functional annotation, we applied Functional Mapping and An-notation (FUMA) [21]. First, functional annotation of all genome-wide significant SNPs (*n* = 772, excluding those not available in FUMA database) showed that significant SNPs were mostly (90.8%) located in intronic/intergenic areas (Supplementary Table S7 and Figure 4b). Those relevant SNPs were also enriched for chromatin states 4 (33.2%) and 5 (40.0%), indicating effects on active transcription (Figure 4a). More important, six SNPs were exonic non-synonymous, which lead to a probably deleterious change in the sequence of the encoded protein (Supplementary Table S8). Five genome-wide significant SNPs (rs10507144, rs3789362, rs4646626, rs6680541, and rs2845871) had a high observed probability of a deleterious variant effect (CADD score [22] ≥ 20). Overall, these results suggest that most GWAS relevant SNPs are located in non-coding regions despite some non-synonymous variants have been found.

**Figure 4:**
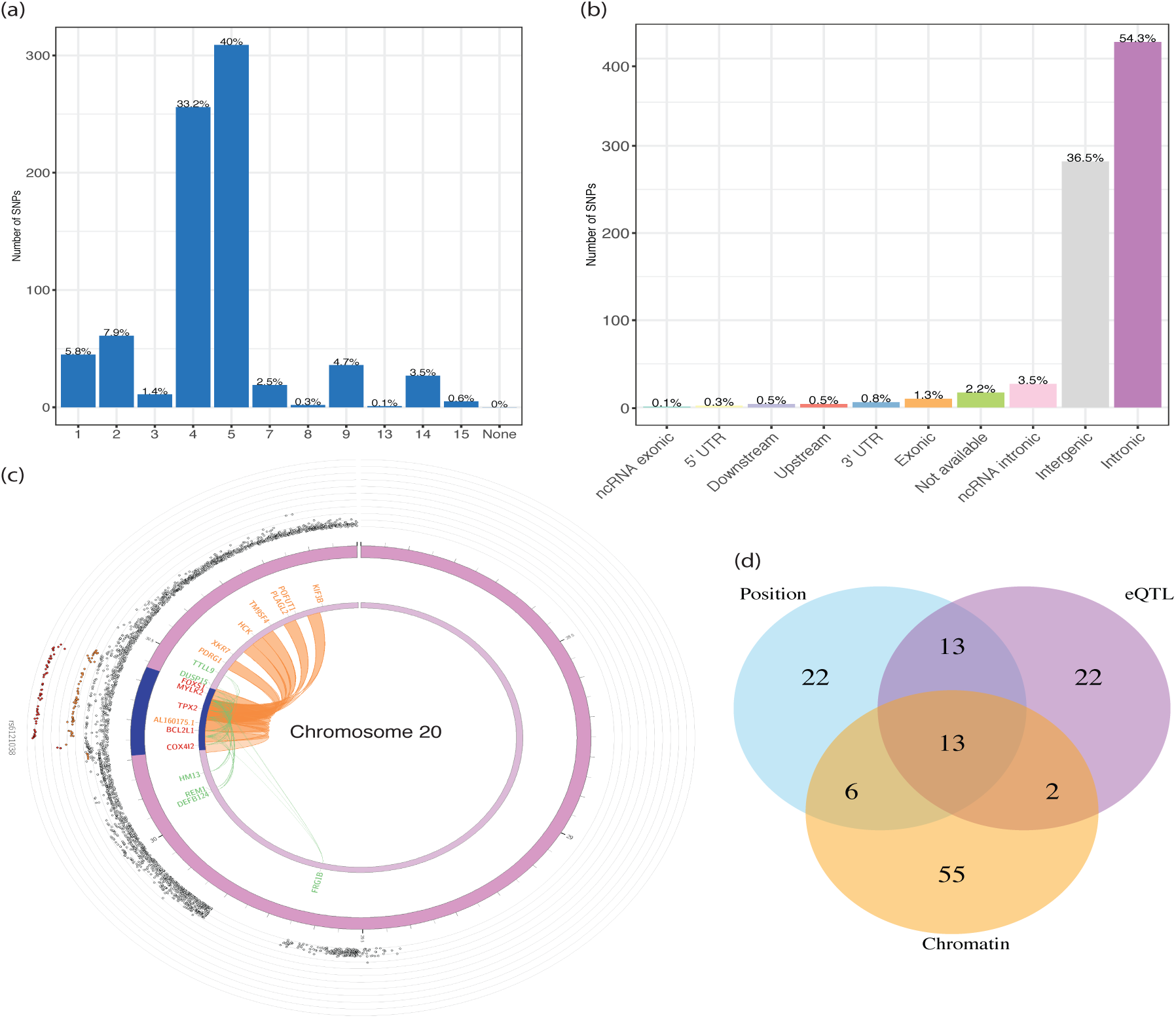
Functional annotation and implications of aMAT results. **a** and **b**, Distribution of (**a**) functional effects and (**b**) minimum chromatin state across 127 tissue/cell types for variants in aMAT identified genomic risk loci. **c**, Zoomed-in Circos plot of chromosome 20. Circos plots show implicated genes by either chromatin interaction (colored orange) or eQTLs (colored green) or both (colored red). The dark blue areas are genomic risk loci identified by aMAT. Chromatin interactions and eQTL associations are colored orange and green, respectively. The most outer layer shows a Manhattan plot, displaying −log_10_(*p*) for SNPs with *p <* 0.05. **d**, Venn diagram of number of linked genes by three different strategies.

Second, we linked the associated variants to genes via three gene-mapping strategies used in FUMA. Figure 4d shows the Venn diagram of the number of mapped genes by three mapping strategies. Positional, eQTL, and chromatin interaction gene mapping strategies linked SNPs to 54, 50, and 76 genes, respectively. This resulted in 133 unique mapped genes (Supplementary Table S9), 13 of which were identified by all three mapping strategies (Supplementary Table S10). The locus (rs6121038) on chromosome 20 is particularly notable. Many genes in this locus (including *FOXS1, MYLK2, TPX2, BCL2L1*, and *COX412*) might interact physically or through eQTL, as indicated by chromatin interaction data in mesenchymal stem cell line and eQTL data (Figure 4c). Therefore, those genes might affect volumetric IDPs via a similar biological mechanism. Six genes (*STX6, EGFR, SMIM19, METTL10, LEMD3*, and *MYLK2*) are of particular interest as they are implicated via eQTL mapping with brain-related tissues. By searching GWAS Catalog [1], we found that three out of six (*METTL10, LEMD3*, and *MYLK2*) genes were reported to be associated with volumetric IDPs. For example, *METTL10* was associated with mean platelet volume [28], total hippocampal volume [29], and dentate gyrus granule cell layer volume [29]. *STX6* was identified by all three mapping strategies and contained a significant SNP (rs3789362) with CADD score higher than 20. *STX6* was reported to be associated with progressive supranuclear palsy (PSP) [30, 31] and the volume of many brain regions (including cerebellum, thalamus, putamen, pallidum, hippocampus, and brainstem) were significantly reduced in PSP compared to control subjects [32]. Future studies might thus consider the potential role of *STX6* in PSP.

Third, the identified genes were enriched in many GWAS Catalog reported volumetric gene sets, including dentate gyrus granule cell layer volume (*P* = 1.46 × 10^−13^), hippocampal subfield CA4 volume (*P* = 1.46 × 10^−13^), and hippocampal subfield CA3 volume (*P* = 5.96 × 10^−12^) (Supplementary Figure S20). We further performed gene-set analysis for tissue expression and biological pathways via FUMA. Two related tissues (Brain Cerebellar Hemisphere and Brain Cerebellum) were nominally associated with the volumetric IDPs set when only correcting the number of tissues being tested (Figure S21).

In summary, aMAT identified several novel and replicable risk loci that have been ignored by standard GWAS analysis. These newly identified loci not only showcase the power of our proposed approach but also deepens our understanding of the genetic basis of brain volumetric measures.

### 3.5 aMAT identifies novel risk loci for four additional IDP sets

We further applied aMAT to four additional IDP sets, which were defined by [11] and represented many IDP classes with significant trait correlations in each grouping (Figure 5). These sets include 206 region area related IDPs (denoted by Area), 208 region thickness related IDPs (denoted by Thickness), 472 structural MRI related IDPs constructed by Freesurfer (denoted by Freesurfer), and 138 T1 ROIs related IDPs (denoted by ROIs). In total, we identified 84 independent risk loci and 97 lead SNPs, 59 of which were ignored by any individual IDP tests at the genome-wide significance level (Table 4, Supplementary Tables S11–S14, and Supplementary Figures S22–S25).

**Table 4:**
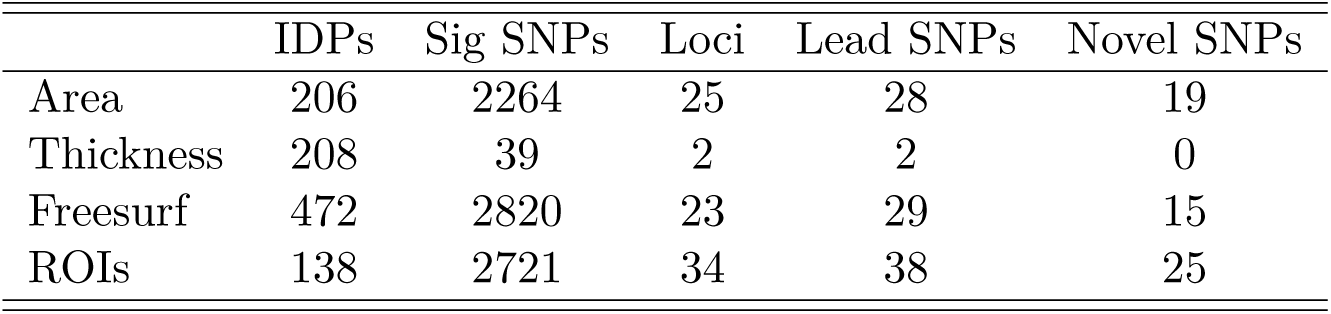
Summary statistics of aMAT for analyzing four additional IDP sets. IDPs, Sig SNPs, Loci, Lead SNPs are the number of IDPs in the set, number of genome-wide significant SNPs identified by aMAT, number of independent risk loci, number of lead SNPs in the identified risk loci, respectively. Novel SNPs is the number of lead SNPs that were ignored by any individual GWAS at genome-wide significance level (5 × 10^−8^).

**Figure 5:**
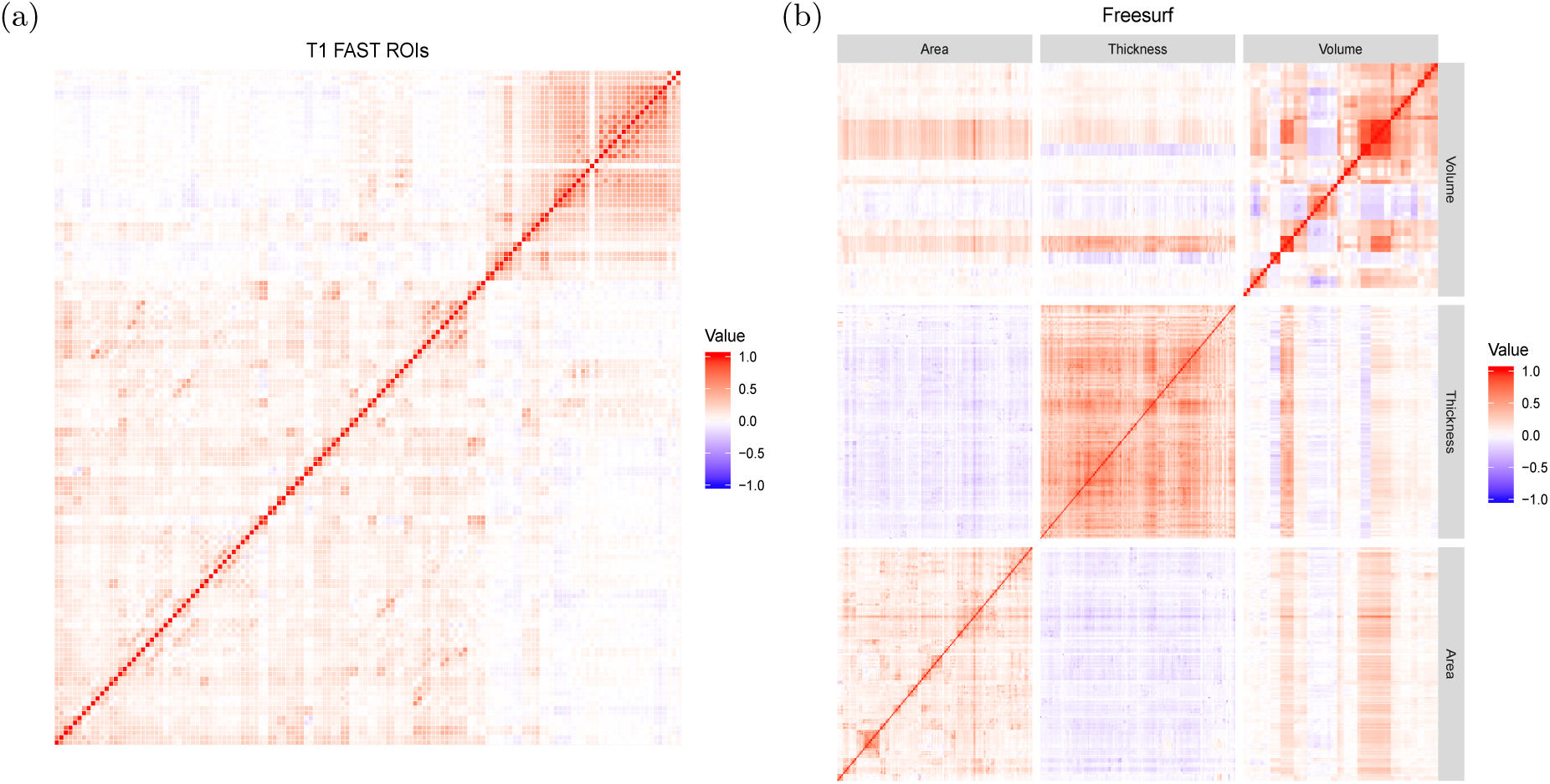
The estimated trait correlation matrix for different sets of IDPs. The subfigure (a) was for T1 FAST region of interests (ROIs) set, which contained 138 IDPs, and subfigure (b) was for Freesurf set, which contained 472 IDPs. Note that the trait correlation estimated here was the correlation of the raw phenotypes conditional on the covariates (e.g., confoundings), which was different from the raw trait correlation provided by Elliott et al. (2018) [11].

### 3.6 aMAT performs better than competing methods in real data applications

We compared aMAT to many competing methods. First, aMAT had a good genomic inflation factor in general (from 0.967 to 1.097) (Supplementary Table S15). In comparison, chi-squared test and Hom test had unstable genomic inflation factors. For example, chi-squared test had a genomic inflation factor of 0.338 for analyzing the Volume IDP set, while Hom test had a genomic inflation factor of 3.102 for analyzing the Freesurf IDP set and 0.755 for analyzing the Area IDP set. SUM had a very good genomic inflation factor, and SSU had a slightly inflated genomic inflation factor (between 1.186 and 1.250). These results are in line with our simulation results.

Next, we compared the statistical power of different tests that have a good genomic inflation factor. Supplementary Figure S26 shows the Venn diagram for the numbers of the significant SNPs and associated loci identified by the tests at the 5×10^−8^ genome-wide significance level for analysing the Volume IDP set. Across the whole genome, aMAT, MAT(50), MAT(1), and SUM identified 801, 671, 3, and 25 significant SNPs, respectively. More important, aMAT, MAT(50), MAT(1), and SUM detected 453, 347, 0, and 18 significant and novel SNPs, respectively; a significant and novel SNP is defined as the genome-wide significant SNP but not detected by any individual GWAS analysis at the 5 × 10^−8^ genome-wide significance level. We further compared the results at the risk region level; a risk region is defined by LDetect [33]. aMAT detected 23 risk regions, while MAT(1) and SUM identified 1 and 3 risk regions, respectively. Each test detected some risk regions that have been ignored by other tests. For example, aMAT and SUM tests identified five and three risk regions that were ignored by other methods, respectively. This illustrates that each test can be more powerful than the others under certain genetic architectures. For analyzing other IDP sets, we obtained similar results (Supplementary Figure S27–S30).

## 4 Discussion

We have introduced aMAT, a multi-trait association test, to conduct jointly analysis of any number of traits. Through simulations and real data analyses, we demonstrated that aMAT could yield well-controlled Type I error rates and achieve high statistical power across a wide range of scenarios. In our empirical application to the Volume set (of 58 volumetric IDPs), we identified 28 lead SNPs located in 24 distinct risk loci, 13 of which were missed by any individual IDP tests at the genomewide significance level 5 × 10^−8^. These identified lead SNPs were well replicated by an independent dataset [27].

aMAT is a computationally efficient method since *p*-values can be calculated analytically. We further improved the computational efficiency by pre-calculating the distribution under the null, which remains the same for all SNPs. For example, by a parallel computing strategy with 100 cores in a standard server (each core has 4 GB memory), aMAT completed calculating *p*-values for multitrait analysis of volumetric set in 2.2 minutes (about 9.97 million SNPs in total, Supplementary Table S16). Of note, because aMAT involves combining results from different tests, it is excepted that aMAT is slightly slower than competing methods. To facilitate data analysis for both statistical and clinical investigators, we have implemented our proposed method aMAT into open-source software.

aMAT is a general framework and can be easily extended to incorporate other multi-trait methods such as MTAG [16], N-GWAMA [34], and HIPO [35]. Specifically, we can treat any other powerful tests as an individual MTA test and then apply aMAT to combine its results to further improve the power. However, since most methods are originally designed for analyzing a few phenotypes jointly and ignore the singularity problems of the trait correlation matrix, some minor adaptions and strict evaluations are needed. Thus, we leave it to our future work. More importantly, we view aMAT to be complementary (rather than superior) to standard GWAS. This is because standard GWAS focus on a single trait and may identify several risk loci that have been ignored by aMAT.

We conclude with two limitations of our proposed method aMAT. First, the optimal choice of Γ is unknown. Of note, the default setting Γ = {1, 10, 30, 50} performs good in both simulations and real data analyses. We further evaluated several different choices of Γ and found that aMAT results were robust to the choice of Γ in many simulation settings (Supplementary Figure S31). However, there is no theoretical guarantee for our default choice of Γ. Second, aMAT applies LDSC to estimate the trait correlation matrix and therefore inherits both benefits and limitations from LDSC. For example, aMAT applies the univariate LDSC to estimate the diagonal elements of the trait correlation matrix, and univariate LDSC is known to be biased for a trait with high (SNP) heritability [36]. We leave these interesting topics to future work.

## Acknowledgements

We thank the associate editor and two reviewers for helpful and insightful comments and the Research Computing Center at Florida State University for providing computing resources.

